# Mutational and bioinformatic analysis of the binding site for the ribonucleotide reductase-specific transcriptional repressor NrdR

**DOI:** 10.64898/2026.05.11.724285

**Authors:** Saher Shahid, Daniel Lundin, Inna Rozman Grinberg, Britt-Marie Sjöberg

## Abstract

The prevalent transcriptional repressor NrdR binds to highly conserved prokaryotic sequences in the promoter regions of operons encoding the essential enzyme ribonucleotide reductase. The NrdR binding sites consist of two partially palindromic 16 bp sequences (NrdR boxes) separated by a 15-16 bp linker sequence. We have assessed the requirement of both boxes for binding, the propensity of different NrdRs to bind to heterologous binding sites, and that the linker sequence is only limited to length and not sequence conservation. As we have observed several deviations from the conserved sequences of the NrdR boxes, we here test the conservation requirements of individual basepairs in the NrdR boxes using a synthetic DNA fragment (Synt DNA) to which the NrdR proteins from the actinomycete *Streptomyces coelicolor* and the gammaproteobacterium *Escherichia coli* bind equally well as to their homologous binding sites. By introducing isolated mutations to Synt DNA and testing the binding capacity of NrdR from *S. coelicolor* and *E. coli* we expand our understanding of what criteria are needed to build a functional binding site for the NrdR repressor.

## Introduction

The *nrdR* gene is found in more than half of all bacterial genomes (Rozman Grinberg *et al*, 2026). It regulates expression of genes encoding the universal and essential ribonucleotide reductase (RNR) enzymes that provide the building blocks for DNA replication and repair in all cells (Hofer *et al*, 2012; Mathews, 2018; Stubbe & Nocera, 2021; Torrents, 2014). The NrdR repressor protein functions by binding to specific sites in the RNR-encoding promoter regions that overlap the RNA polymerase binding site. It has two binding sites for adenine nucleotides, an ‘inner’ and an ‘outer’ site. When ATP levels are high both sites are occupied by ATP and the NrdR protein adopts higher oligomeric forms that are unable to bind to DNA. However, when dNTP levels increase as a result of halted DNA synthesis, the outer site is invaded by dATP, inducing a tetrameric form that binds to DNA and inhibits expression of RNR-encoding genes.

The NrdR binding site consists of two partially palindromic 16 bp NrdR boxes (box 1 and box 2) at a distance of 15-16 bp. The partially conserved palindromic sequence of an NrdR box was initially predicted from a small number of bacterial genomes (Rodionov & Gelfand, 2005). Later a bioinformatic analysis of species-representative genomes revealed the consensus sequence for the NrdR boxes (Fig. 1), where some positions in each box are highly conserved (Rozman Grinberg *et al*, 2022). A mutational study in *Streptomyces coelicolor* demonstrated that C4 and G13 were crucial entities of the binding site (Rozman Grinberg *et al*., 2022), and recent *in vivo* experiments established that a correct spacing between the two boxes are crucial for high-affinity binding (Shahid *et al*, 2025).

**Figure 1.**
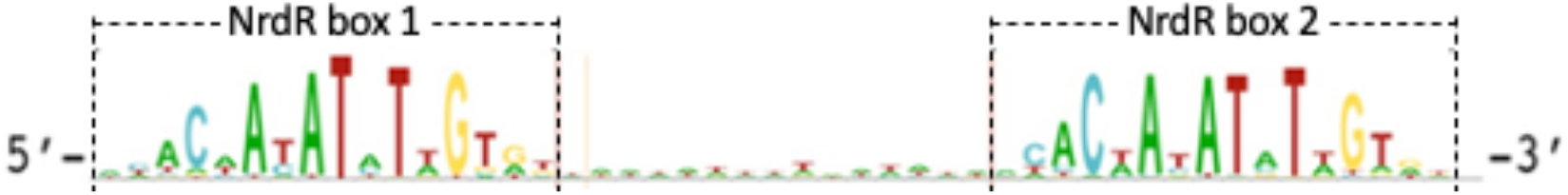
Logo of NrdR boxes from 935 sequences. (Rozman Grinberg *et al*., 2022).

NrdR’s function has primarily been experimentally studied in *Streptomyces coelicolor* with two RNR-encoding operons (*nrdAB* and *nrdRJ*), *Escherichia coli* with three RNR-encoding operons (*nrdAB, nrdHIEF*, and *nrdDG*), and *Pseudomonas aeruginosa* with three RNR-encoding genomes (*nrdAB, nrdJab*, and *nrdDG*) (Crespo *et al*, 2015; Grinberg *et al*, 2006; Pedraz *et al*, 2026; Rozman Grinberg *et al*, 2025; Rozman Grinberg *et al*., 2022). Since each of these three genomes display in their NrdR binding sites discrepancies from the consensus sequence of NrdR boxes we have performed an extended mutational study to understand which nucleotides in the NrdR boxes are critical for binding to DNA. In this study, the NrdR protein from *E. coli* (EcoNrdR) and *S. coelicolor* (ScoNrdR) were analyzed for their ability to bind to wild type and mutated NrdR boxes using microscale thermophoresis (MST).

## Results and Discussion

### Heterologous binding of NrdR to NrdR boxes

To establish whether NrdR from *E. coli* (EcoNrdR) can bind *S. coelicolor* NrdR boxes and vice versa we compared binding of the two NrdRs to all five promoter regions. Binding of *S. coelicolor* NrdR (ScoNrdR) to *E. coli* RNR promoters were comparable to binding to its own promoter regions and resulted in binding affinities between 56 and 169 nM (Table 1, Supplementary fig. 1A). Similarly, EcoNrdR bound to *S. coelicolor* promoter regions with affinities comparable to affinities to its own promoter regions and had affinities of 126 and 347 nM (Table 1, Supplementary fig. 1B). These results indicate that NrdR from one organism can bind to NrdR boxes from other organisms due to similarity between the boxes as well as the proteins. Earlier studies have for instance demonstrated a striking structural similarity of the DNA-binding tetramers of ScoNrdR and EcoNrdR (Rozman Grinberg *et al*., 2025; Rozman Grinberg *et al*., 2022).

**Table 1.**
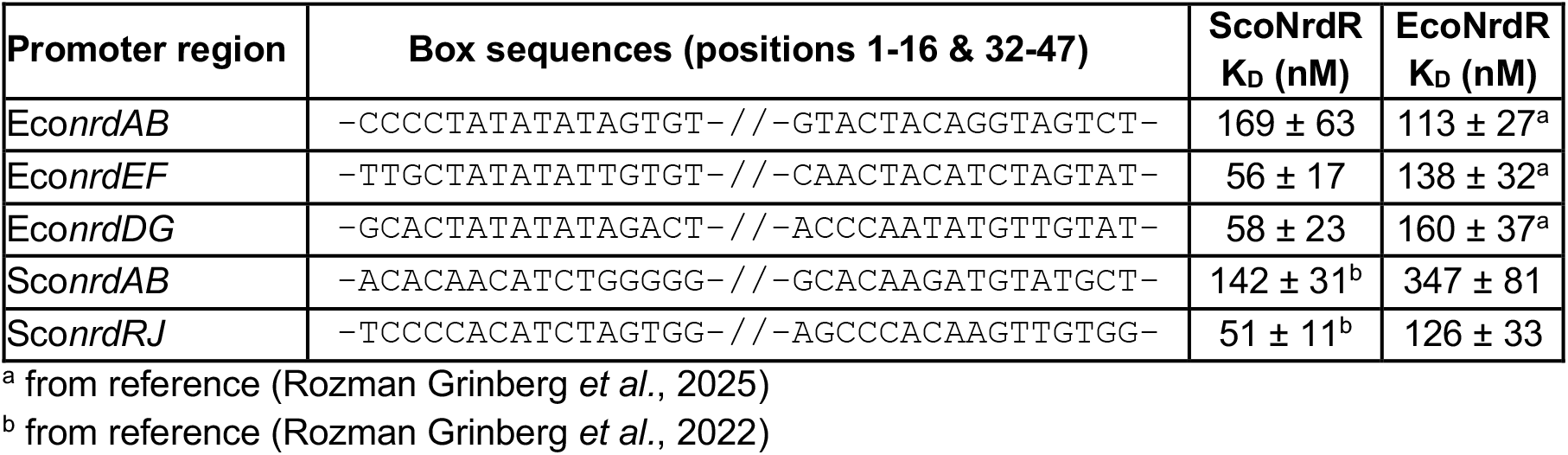
Heterologous binding of NrdR to RNR promoter regions.

### Design of a synthetic NrdR binding site

Initially, we intended to use short DNA fragments containing a single NrdR box with five flanking base pairs, and to mutate the box nucleotides one by one to test how each individual base pair affects binding affinity. However, the NrdR protein requires two NrdR boxes to bind to DNA, as shown in figure 2 for the lack of binding of *E. coli* NrdR to isolated NrdR box 1 or NrdR box 2.

**Figure 2:**
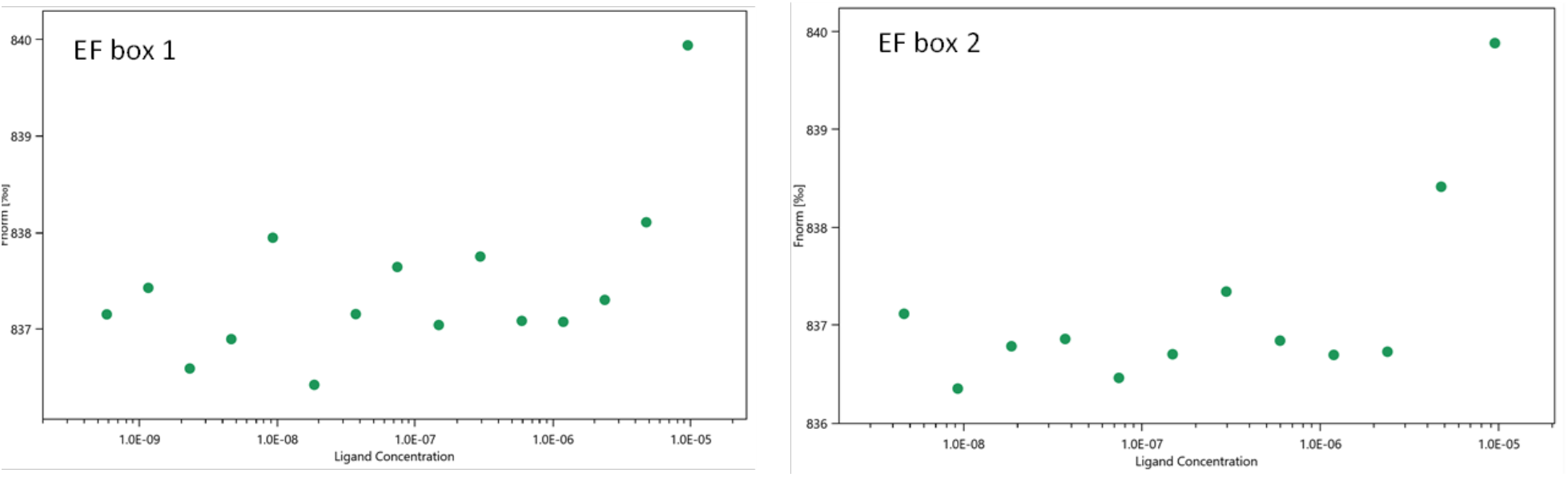
Binding of EcoNrdR to individual boxes of the *E. coli nrdEF* promoter.

We therefore designed a synthetic NrdR binding site denoted ‘Synt DNA’, consisting of two identical NrdR boxes connected via a 15-bp linker corresponding to the linker region of the *S. coelicolor nrdRJ* promoter (Fig. 3). The Synt DNA binding site is flanked by 5 bp at each end. Both ScoNrdR and EcoNrdR had high affinity to the 57-bp Synt DNA with K_D_s comparable to binding to their wild type promoters (Fig. 3 and Table 1).

**Figure 3.**
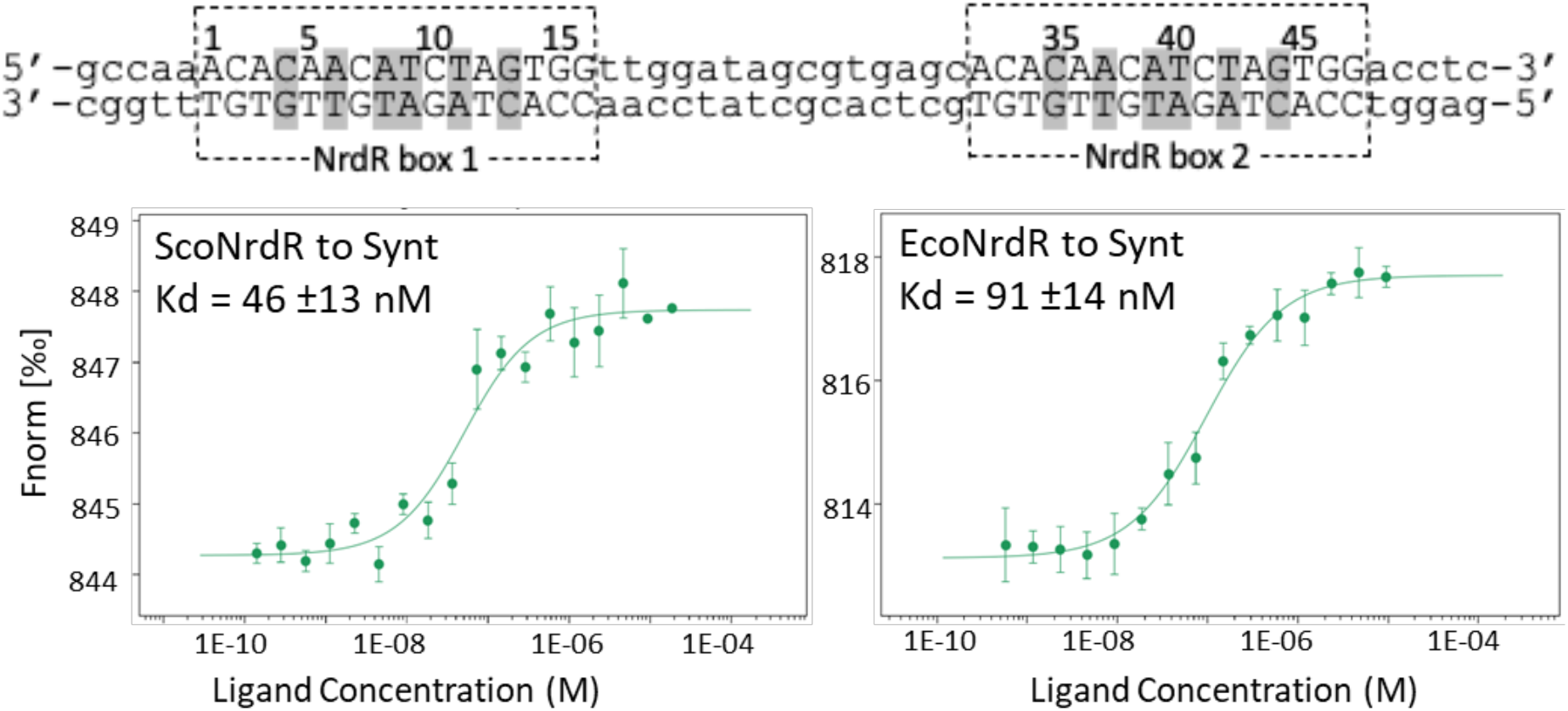
Binding of ScoNrdR and EcoNrdR to a synthetic NrdR box region based on the NrdR box logo sequence. A) Sequence of a 57 bp synthetic DNA fragment (Synt DNA) with two identical NrdR boxes separated by a 15 bp linker sequence with numbering starting at NrdR box 1. NrdR boxes are in capital letters, highly conserved NrdR box positions shaded in gray, and flanking regions and the linker sequence in lowercase letters. B) MST binding curves.

The linker region between box 1 and box 2 in natural binding sites is not conserved. Changing the linker region between the synthetic NrdR boxes from the *S. coelicolor nrdRJ* linker to the *S. coelicolor nrdAB* linker, which have only 13% identity, had no effect on the binding affinity of ScoNrdR or EcoNrdR (Table 2, Supplementary fig. 2). This finding fits well with our earlier observation that NrdR proteins from four different species had the capacity to bind to linker regions corresponding to the predominant length of 15-16 bp as well as a linker length of 26-27 bp, i.e. adding another DNA helix turn to the distance between NrdR box 1 and 2 (Shahid *et al*., 2025).

**Table 2.**
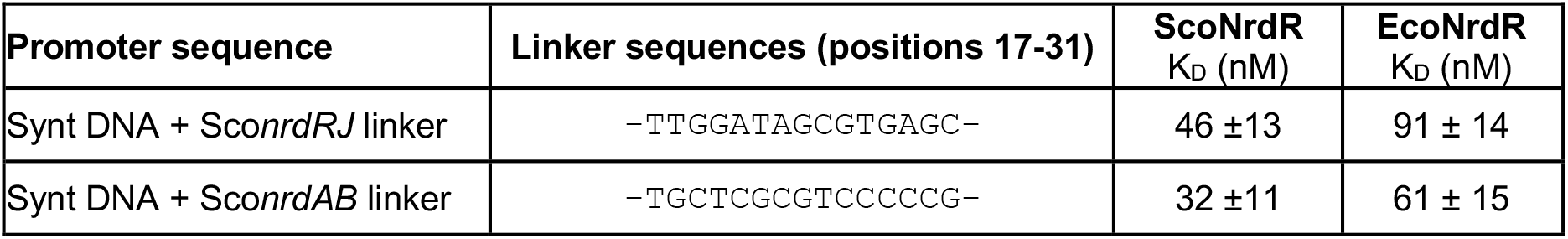
Effect of linker sequence on binding affinity of EcoNrdR and ScoNrdR.

### Mutations affecting highly conserved NrdR box positions

The consensus NrdR box is a partially palindromic 16 bp sequence in which 6 positions are highly conserved: C4, A6, A8, T9, T11, G13 (Fig. 1). We previously showed that C4 and G13 in both NrdR boxes are critical for ScoNrdR binding to its native DNA sequence (Rozman Grinberg *et al*., 2022). Inversion or mutation of these nucleotides resulted in significant loss of affinity. To confirm this for EcoNrdR we introduced the single mutation G44C in the native *E. coli nrdEF* NrdR box 2, which completely abolished binding (Table 3, Supplementary fig. 3B). On the other hand, a comparative study of NrdR boxes in 6 different *Mycobacteriaceae* species predicted that box 1 in the promoter region of the *nrdHIE* operon had a G in position 4, whereas box 1 had a C4 in the promoter region of the *nrdF2* gene (Mowa *et al*, 2009). We are not aware of any experimental studies to substantiate these predictions.

**Table 3.**
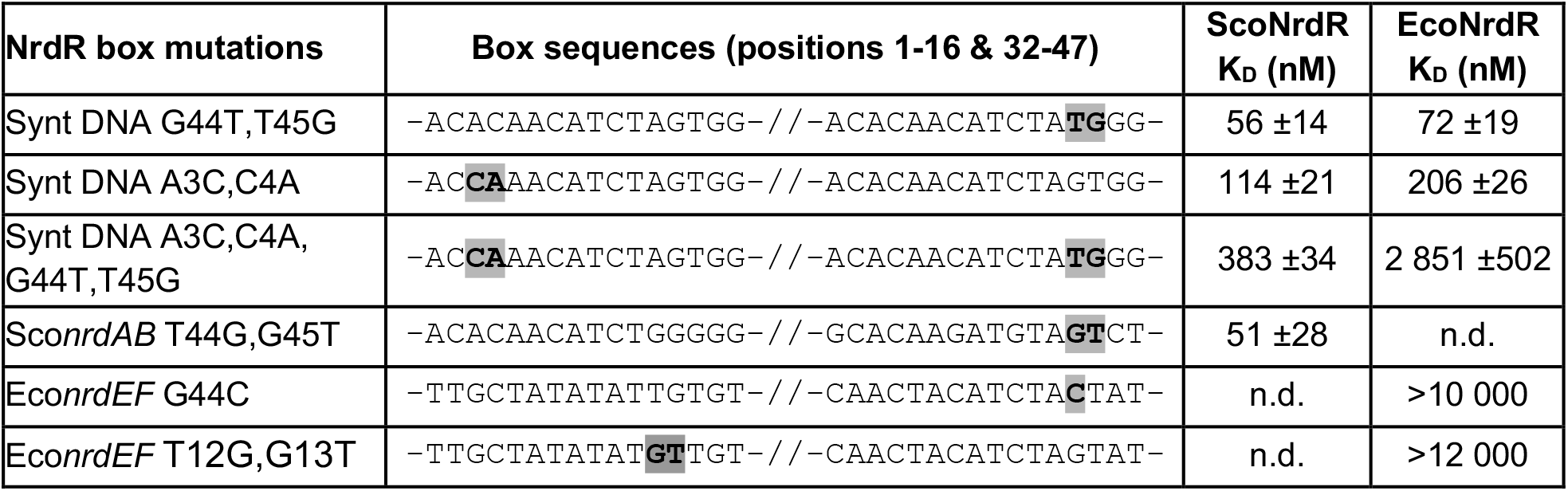
**Binding of ScoNrdR and EcoNrdR to mutated Synt DNA fragments and mutated RNR promoter regions.** (n.d., not determined)

The distance between C and G is usually 8 bp, but the *S. coelicolor nrdAB* promoter has a distance of 9 bp (Table 1). We investigated whether these conserved Cs and/or Gs could be moved one position in box 1, box 2 or both boxes of the Synt DNA sequence such that there are 9 bp between C and G. Mutations G44T plus T45G in box 2 resulted in high affinity binding of both EcoNrdR and ScoNrdR (Table 3, Supplementary fig. 3). The same was true when C4 was moved one position with mutations A3C plus C4A in box 1 of the Synt DNA sequence. However, moving C4 and G44 at the same time (A3C, C4A; G44T, T45G), so that there was 9 bp distance between the conserved C and G in both boxes, resulted in significant loss of binding (Table 3, Supplementary fig. 3). To explore whether a 7 bp distance is acceptable between C and G in a NrdR box, we mutated the first NrdR box in the *E. coli nrdEF* promoter (T12G,G13T). This mutant lost the ability to bind NrdR (Table 3, Supplementary fig. 3B) suggesting that a distance of at least 8 bp is required between C and G.

A6 and T11 are highly conserved. Mutating A6 to C in both NrdR boxes (A6C plus A37C) resulted in more than 20-fold decrease in binding affinity for both EcoNrdR and ScoNrdR (Table 4, Supplementary fig. 4). Moreover, mutation of both A6 and T11 in both NrdR boxes (A6C, T11G, A37C, T42G) abolished the binding completely (Table 4, Supplementary fig. 4). For EcoNrdR mutating A6 to G in both NrdR boxes decreased binding affinity 10-fold, whereas binding of ScoNrdR was decreased 4-fold (Table 4, Supplementary fig. 4), suggesting that a GC base pair at this position significantly impaired NrdR binding. On the other hand, mutating A6 to T in both boxes didn’t have an effect on EcoNrdR binding, but still decreased ScoNrdR binding 3-fold (Table 4, Supplementary fig. 4). One additional aspect of the conservation of positions 6 and 11 in both boxes (6/11 and 37/42) could be the partially palindromic nature of each box, a feature that may be important *in vivo*, but which will plausibly not affect K_D_s measured *in vitro* using short DNA fragments.

**Table 4.**
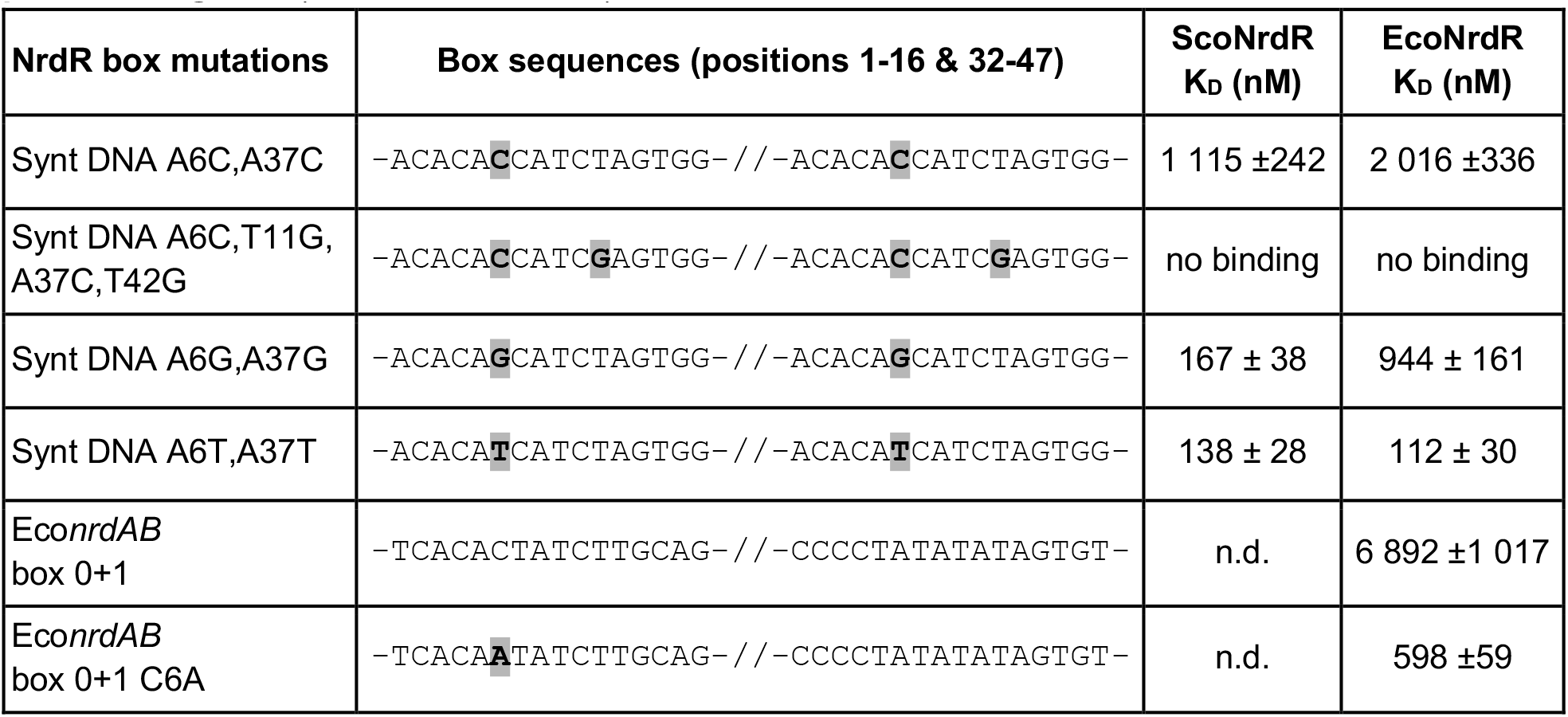
**Binding of ScoNrdR and EcoNrdR to mutated Synt DNA fragments and *E. coli* RNR promoter regions.** (n.d., not determined)

Earlier studies have annotated a different box pair in the *E. coli nrdAB* promoter consisting of box 1 and an upstream box covering the -10 region and the transcriptional start site (Rodionov & Gelfand, 2005; Torrents *et al*, 2007). However, we showed recently that this upstream box, denoted box 0, is not recognised by EcoNrdR (Rozman Grinberg *et al*., 2025). We tested if the lack of binding to box 0 is that position 6 in this case is a C instead of A, and found that the mutation C6A in box 0 resulted in 10 times higher affinity of EcoNrdR (Table 4, Supplementary fig. 4).

### The conserved middle AT sequence

The consensus NrdR box sequence predicts a middle conserved AT sequence (Rodionov & Gelfand, 2005; Rozman Grinberg *et al*., 2022). Yet, box 2 in the *S. coelicolor nrdRJ* promoter region has a middle AA sequence, and box 2 in the *E. coli nrdAB* promoter region has a middle AG sequence (Table 1). We therefore performed a bioinformatic analysis of close to 60 000 NrdR binding motifs in species-representative genomes from all bacterial species, and as expected found that a middle AT sequence is predominant (Figure 4). A middle AA sequence is 6 times less frequent, and middle TT and middle TA sequences are even less frequent.

**Figure 4.**
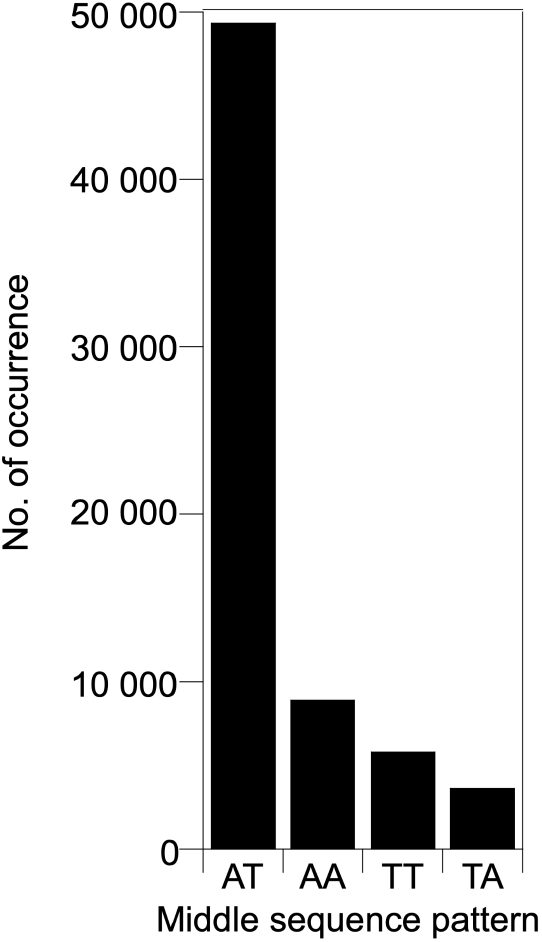
Frequencies of middle nucleotide sequence in NrdR boxes.

We used our Synt DNA to experimentally check the importance of conserved A8 and T9. Mutation T9A and T40A resulting in a middle AA in one or both NrdR boxes respectively and mutation A8T and A39T resulting in middle TT in both boxes still had preserved binding affinity (Table 5, Supplementary fig. 5). As the *S. coelicolor nrdRJ* promoter NrdR box 2 contains a middle AA sequence, we also tested whether ScoNrdR would tolerate a middle AA also in box 1 and introduced a T9A mutation in its natural NrdR binding site. Sco*nrdRJ* T9A had a comparable K_D_ of 83 nM to the K_D_ of 51 nM for ScoNrdR binding to its natural Sco*nrdRJ* binding site, showing that ScoNrdR can recognize a binding site with a middle AA sequence in both NrdR boxes (Table 5, Supplementary fig. 5A).

**Table 5.**
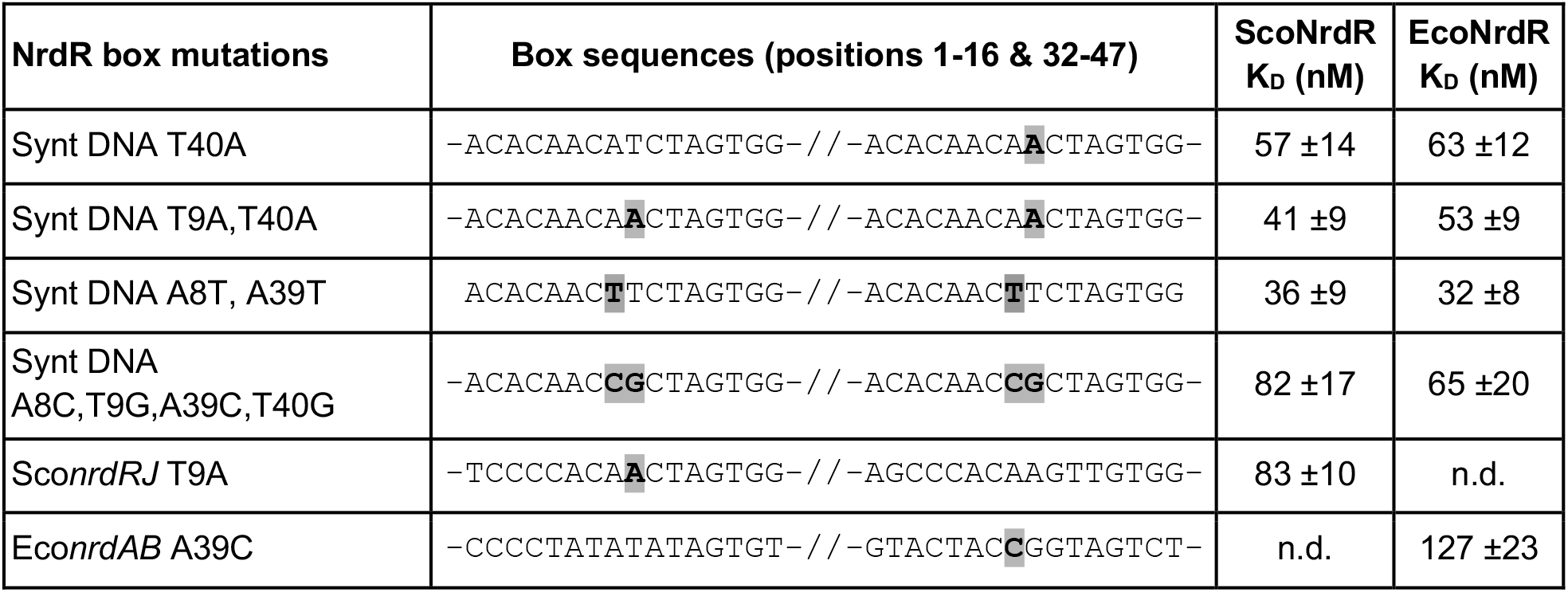
**Binding of ScoNrdR and EcoNrdR to mutated Synt DNA fragments and mutated RNR promoter regions.** (n.d., not determined)

The seemingly drastic mutations A8C plus T9G in both NrdR boxes had only marginal effect on binding affinity for both ScoNrdR and EcoNrdR (Table 5, Supplementary fig. 5). To confirm this surprising result, we also introduced an A39C mutation in the natural NrdR binding site of the *E. coli nrdAB* promoter region, which has a middle AG sequence in box 2, changing its box 2 to a middle CG sequence. EcoNrdR binds to the Eco*nrdAB* A39C NrdR binding site with similar affinity, K_D_ of 127 nM compared to K_D_ of 113 nM (Table 5, Supplementary fig. 5B).

#### Total AT content in the central part of an NrdR box

High resolution structures of ScoNrdR and EcoNrdR bound to DNA show that the dsDNA helix adopts sharp bends at each NrdR box (Rozman Grinberg *et al*., 2025; Rozman Grinberg *et al*., 2022). This was primarily ascribed to the predominant middle AT sequence of the NrdR box. As both bioinformatics and experimental studies (Fig. 4, Table 5) (Crespo *et al*., 2015; Grinberg *et al*., 2006; Pedraz *et al*., 2026; Rozman Grinberg *et al*., 2025; Rozman Grinberg *et al*., 2022) show that a middle AT sequence is not conserved, we checked the total AT content in the central part of NrdR box sequences in a few bacterial species with different genomic GC contents. The content of AT basepairs spans from 63% to 100% between the conserved positions C4 and G13 in native NrdR boxes, and the fraction of AT basepairs does not seem to correlate with the genomic GC content of the species (Table 6). The surprising discrepancy between a consensus box sequence and our finding that a middle AT sequence is not necessarily required conceivably relates to the fact that the rare occurrences of other basepairs in positions 8 and 9 are compensated by adjacent high content of AT basepairs still allowing the sharp bend of DNA required for binding of the NrdR protein (Rozman Grinberg *et al*., 2025; Rozman Grinberg *et al*., 2022). Despite the substantial sequence variation in NrdR boxes between bacterial species, the overall ability of NrdR to recognize and bind these promoter sites is preserved. This suggests that the evolution of NrdR binding sites involves DNA flexibility and spacing in addition to some key conserved residues to achieve optimal interaction.

**Table 6.** AT content in NrdR boxes between positions C4 and G13 for different RNR promoter regions in species with different genomic GC content.

| Species | GC content (%) | Fraction AT basepairs |  |  |  |
| --- | --- | --- | --- | --- | --- |
|  |  | *nrdAB* | *nrdEF* | *nrdJ* | *nrdDG* |
| *Listeria monocytogenes*<sup>a</sup> | 38 | 0.88; 0.94 |  |  | 0.88 |
| *Streptococcus thermophilus* | 39.1 |  | 1.0 |  | 1.0 |
| *Streptococcus pneumoniae* | 39.7 |  | 0.96 |  | 0.88 |
| *Chlamydia trachomatis* | 41.3 | 0.88 |  |  |  |
| *Escherichia coli*<sup>b</sup> | 50.8 | (0.88), 0.81 | 0.88 |  | 0.94 |
| *Salmonella typhimurium* | 52.2 | 0.88 | 0.75 |  | 0.88 |
| *Pseudomonas aeruginosa*<sup>c</sup> | 65-67 | 0.88 |  | 0.81 | 0.75 |
| *Thermotoga maritima*<sup>d</sup> | 67-69 |  |  | 0.79 | 1.0 |
| *Streptomyces coelicolor*<sup>e</sup> | 72 | 0.71 |  | 0.63 |  |

<sup>a</sup>Lm encodes two pairs of NrdR boxes 113 bp apart in the *nrdAB* promoter region.
<sup>b</sup>Ec MG1655 encodes an NrdR box 0 in the *nrdAB* promoter region, which does not bind NrdR.
<sup>c</sup>The Pa genome encodes a split *nrdJ* denoted *nrdJab*.
<sup>d</sup>The Tm *nrdJ* promoter region has three NrdR boxes with a 14 bp linker in both cases.
<sup>e</sup>The second NrdR box in the Sc *nrdAB* promoter region has a T in position 13 (instead of G) and a G in position 14.

### Mutations affecting non-conserved NrdR box positions

Mutating A5 to C in both boxes (A5C plus A36C) had no effect on binding affinity (Table 7, Supplementary fig. 6). However, introducing mutations in both A5 and A12 in both boxes (A5C, A12G, A36C, A43G) resulted in a binding affinity that was 10 times lower for ScoNrdR and 5 times lower for EcoNrdR. The native *S. coelicolor nrdRJ* promoter contains C in position 5 in both NrdR boxes, but either A or T in position 12. Mutation of A12 to G plus T43 to G in the *S. coelicolor nrdRJ* promoter resulted in 10 times decreased affinity (Supplementary fig. 6A) confirming the results with the Synt DNA.

**Table 7.** **Binding of ScoNrdR and EcoNrdR to mutated Synt DNA fragments and mutated *S. coelicolor* RNR promoter regions.** (n.d., not determined)

| NrdR box mutations | Box sequences (positions 1-16 & 32-47) | ScoNrdR<br>K<sub>D</sub> (nM) | EcoNrdR<br>K<sub>D</sub> (nM) |
| --- | --- | --- | --- |
| Synt DNA A5C,A36C | -ACAC<sup>C</sup>ACATCTAGTGG-// -ACAC<sup>C</sup>ACATCTAGTGG- | 62 ±12 | 93 ±21 |
| Synt DNA<br>A5C,A12G,A36C,A43G | -ACAC<sup>C</sup>ACATCT<sup>G</sup>GTGG-// -ACAC<sup>C</sup>ACATCT<sup>G</sup>GTGG- | 447 ±106 | 495 ±107 |
| SconrdRJ A12G,T43G | -TCCCCACATCT<sup>G</sup>GTGG-// -AGCCCACAAGT<sup>G</sup>GTGG- | 597 ±121 | n.d. |
| Synt DNA<br>C7T,C10A,C38T,C41A | -ACACAA<sup>T</sup><sup>A</sup>TATAGTGG-// -ACACAA<sup>T</sup><sup>A</sup>TATAGTGG- | 45 ±10 | 41 ±15 |
| Synt DNA A3C,A34C | -AC<sup>C</sup>CAACATCTAGTGG-// -AC<sup>C</sup>CAACATCTAGTGG- | 46 ±8 | 37 ±10 |
| Synt DNA T14G,T45G | -ACACAACATCTAG<sup>G</sup>GG-// -ACACAACATCTAG<sup>G</sup>GG- | 46 ±11 | 53 ±10 |
| Synt DNA<br>A3C,T14G,A34C,T45G | -AC<sup>C</sup>CAACATCTAG<sup>G</sup>GG-// -AC<sup>C</sup>CAACATCTAG<sup>G</sup>GG- | 33 ±7 | 95 ±12 |

Mutating the non-conserved C7 and C10 in both NrdR boxes (C7A, C10T, C38T, C41A) did not affect the binding affinity of neither ScoNrdR nor EcoNrdR (Table 7, Supplementary fig. 7). Moreover, a duplication of *E. coli nrdEF* box 1 (with A7 and T10) or box 2 (with C38 and C41) had no effect on binding affinity (Shahid *et al*., 2025), corroborating that positions 7 and 10 in the NrdR boxes are not conserved.

Mutations in position 3 and 14 towards the edges of the boxes, separately and together, had as expected very little effect on affinity (Table 7, Supplementary fig. 8). These results exclude the possibility that a C and/or G next to C3 and/or G13 (as exists in *S. coelicolor* promoters) improves recognition of the box by NrdR.

## Concluding remarks

In this work, we performed a systematic mutational analysis of NrdR boxes and demonstrated how specific alterations influence the binding strength of *S. coelicolor* and *E. coli* NrdR proteins. The key findings are summarized schematically in Scheme 1.

**Scheme 1.**
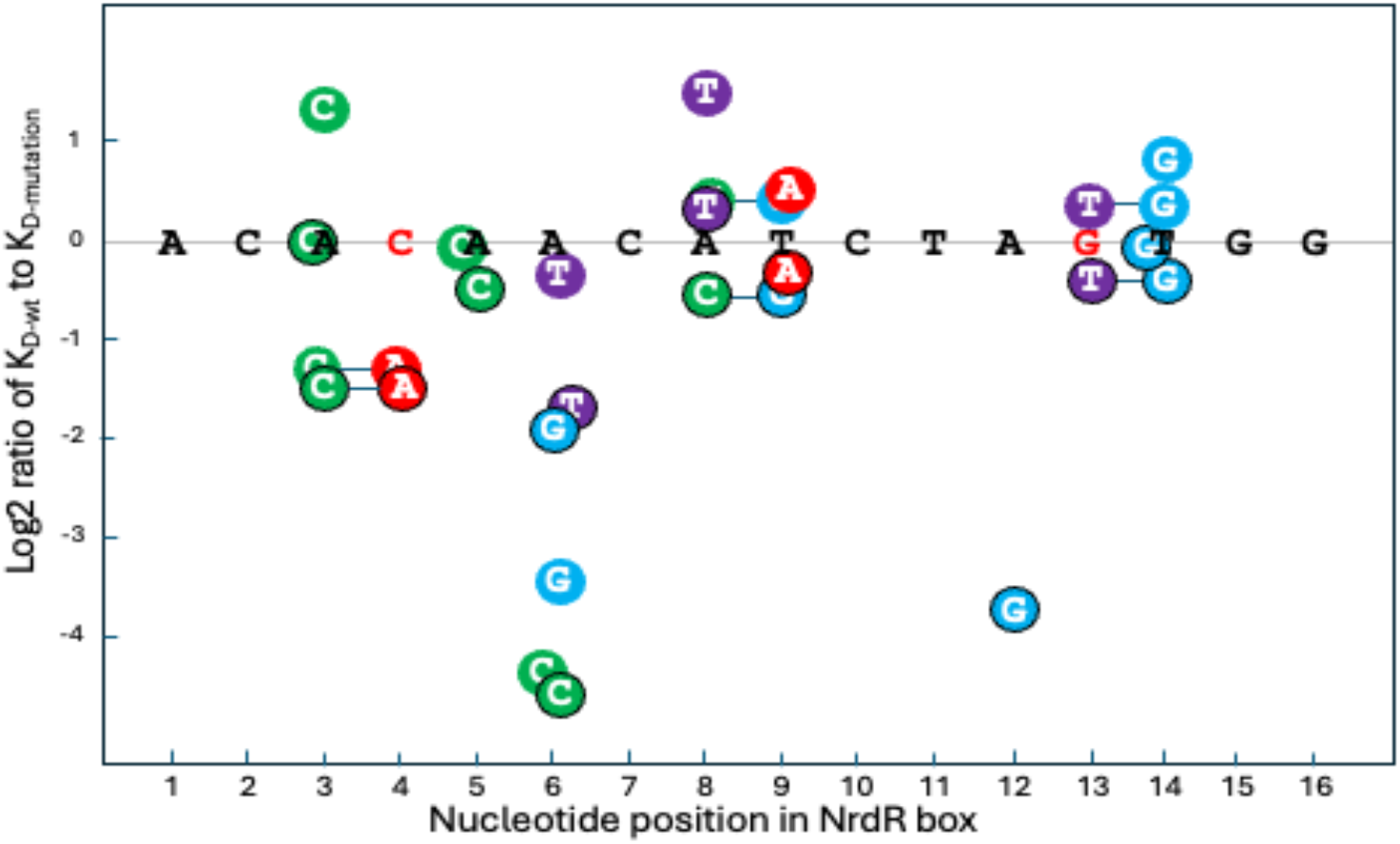
Summary of single and double mutation effects on binding of EcoNrdR and ScoNrdR to Synt DNA fragments. The Synt DNA sequence is shown in black, with essential C4 and G13 in red. Mutation effects with black contours refer to SocNrdR binding, and without contours to EcoNrdR binding. Double mutations are connected via a bar. Positive log2 values indicate mutations with stronger binding than wt, and negative log2 values mutations with worse binding compared to the non-mutated Synt DNA. Mutation values close to wt values are slightly offset from their sequence positions for increased clarity.

A bioinformatic study of species-representative genomes showed that positions C4, A6, A8, T9, T11 and G13 are highly conserved in each of the two NrdR boxes constituting the binding site for the prevalent prokaryotic transcriptional repressor NrdR (Rozman Grinberg *et al*., 2022). At the same time part of the NrdR binding site is also a binding site for sigma factors, RNA transcriptase subunits and/or other regulatory factors, which may explain why some deviations from these conserved positions occur. Based on this experimental mutation study of the NrdR boxes we can now summarize the following observations:

- NrdR binding requires a dual-box architecture where both boxes are necessary for efficient DNA binding.
- C4 and G13 is critical, and we showed earlier that these bases bind to residues Asp15 and Arg17 in *S. coelicolor* NrdR (Rozman Grinberg *et al*., 2022), but a distance of 9 bp rather than 8 bp between C and G is tolerated in one box but not in both.
- A6 and T11 seem critical, but no specific interaction with the NrdR protein has been identified. Plausibly the partially palindromic structure of 6 and 11 is important *in vivo*.
- A8 and T9 are not absolutely critical for binding. Rather there is a lower limit of 63% AT base pairs needed between positions C4 and G13 to allow for the bend of dsDNA in the palindromic NrdR box.

Our results suggest that NrdR recognizes DNA through a combination of specific base interactions and overall shape and spacing of the DNA. This study will enable more precise identification of NrdR binding sites across bacterial genomes to support the exploration of NrdR and its RNR regulons as potential targets for antimicrobial development in clinically relevant pathogens.

## Materials and Methods

### Bioinformatic analysis

After identifying potential RNR operons – genes annotated as RNRs by Prokka (v. 1.14.6, (Seemann, 2014)), separated by no more than 300 nucleotides – in all species-representative bacterial genomes in the GTDB (release R08-RS214, (Parks *et al*, 2018)) 500 nucleotides upstream of each operon were extracted. Each fragment was searched with regular expression for a pair of NrdR boxes separated by a spacer of no more than 50 nucleotides (“C.[AT].[AT][AT].[AT].G.{1,50}?C.[AT].[AT][AT].[AT].G”). Included in the search were sequences matching spacers with 20 to 23 nucleotides (corresponding to a box distance of 14-17 bp; 25934 occurrences) and spacers with 30 to 33 nucleotides (corresponding to a box distance of 24-27 bp; 7218 occurrences).

### Synthetic sequence and mutants

Cy5-labelled sense strands and unlabelled antisense strands were generally 57 bp long and purchased from Sigma-Aldrich. For simplicity only Synt DNA wt is shown (Table 8). To elucidate binding of NrdR to individual boxes, shorter oligonucleotides of 26 bases were ordered, containing either individual NrdR box 1 of *nrdEF* RNR promoter or box 2 of *nrdEF* RNR promoter (Table 8).

**Table 8.**
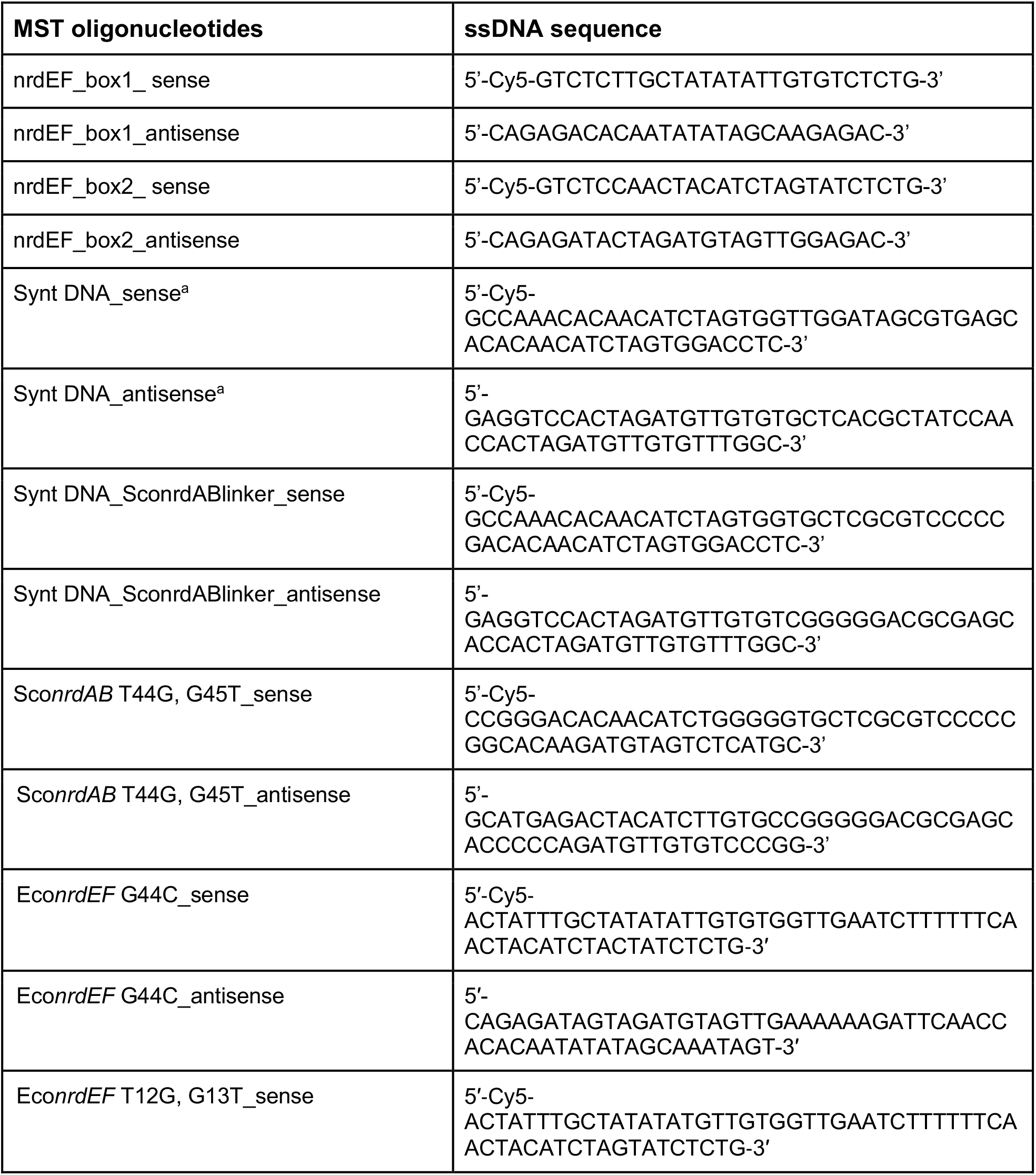
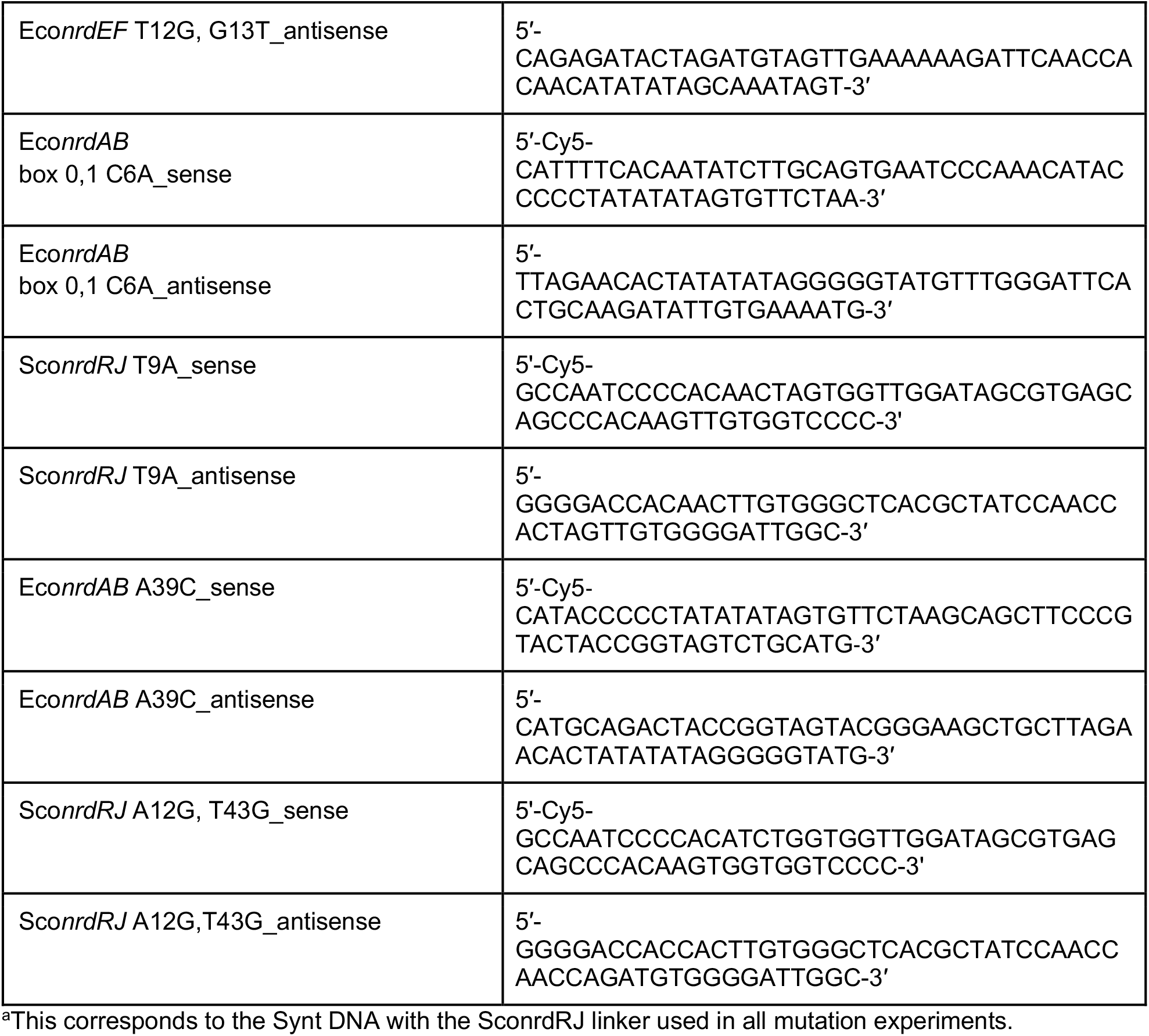
List of oligonucleotides used in MST experiments. Additional oligonucleotides used in this study have been published (Rozman Grinberg *et al*., 2025; Rozman Grinberg *et al*., 2022; Shahid *et al*., 2025).

### Plasmids and reagents

The *nrdR* gene from *S. coelicolor* strain M145 and from *E. coli* K-12 strain MG1655 were amplified and cloned into the pET30a(+) vector-Novagen (Merck KGaA, Darmstadt, Germany) as described previously (Grinberg *et al*., 2006; Torrents *et al*., 2007), resulting in pET30a(+)::Sco*nrdR* and pET30a(+)::Eco*nrdR* constructs expressing wild-type NrdR proteins with C-terminal hexahistidine (His-) tags. ATP and dATP solutions (100 mM, pH 7.0) were purchased from Thermofisher Scientific.

### Protein expression

*E. coli* BL21(DE3) were transformed with the pET30a(+) vector harboring the *EcoNrdR*/*ScoNrdR* gene. A single colony of the transformed cells was used to inoculate the LB medium having kanamycin (50 μg/ml) and grown overnight at 37 °C. This overnight culture was used to inoculate LB medium until the OD_600_ of 0.1 and the cells were allowed to grow under vigorous shaking at 37 °C. When the OD_600_ of 0.8-0.9 was achieved, the culture media was cooled down with ice followed by addition of 0.5 mM IPTG and 0.1 mM Zn(CH_3_CO_2_)_2_. Then the cultures were grown overnight at 20 °C and the cells expressing the recombinant protein were harvested by centrifugation.

### Purification of ScoNrdR

The bacterial cell pellet, expressing ScoNrdR, was resuspended in the lysis buffer (0.05 M Tris-HCl (pH 8.0 at 4°C), 0.3 M NaCl, 10% glycerol, 10 mM Imidazole and 2 mM DTT). Phenylmethylsulfonyl fluoride (PMSF) was added to 1 mM to the cell suspension which was then sonicated in an ultrasonic processor (Misonics) until a clear lysate was obtained. The lysate was centrifuged at 18,000 x g at 4°C for 45 min and the supernatant was used for protein purification.

For purification, the supernatant was loaded onto HisTrap FF Ni-Sepharose column (Cytiva) which was pre-equilibrated with the lysis buffer. After loading the protein, the column was extensively washed with 10 and 60 mM Imidazole. The His-tagged ScoNrdR was eluted with the elution buffer (0.05 M Tris-Cl (pH 8.0 at 4°C), 0.3 M NaCl, 10% glycerol, 0.5 M Imidazole and 2 mM DTT). The purified protein was then desalted using HiPrep 26/10 Desalting column (Cytiva) equilibrated with the ScoNrdR storage buffer (0.05 M Tris-Cl (pH 8.0 at 4°C), 0.3 M NaCl, 10% glycerol, and 1 mM TCEP) and stored at -80°C.

### Purification of EcoNrdR

For purification of EcoNrdR, the bacterial cell pellet, expressing EcoNrdR, was resuspended in the lysis buffer (0.05 M Tris-Cl (pH 8.5 at 4°C), 0.3 M NaCl, 10 mM Imidazole and 2 mM DTT). Phenylmethylsulfonyl fluoride (PMSF) was added to 1 mM to the cell suspension which was then sonicated in an ultrasonic processor (Misonics) until a clear lysate was obtained. The lysate was centrifuged at 18,000 x g at 4°C for 45 min and the supernatant was used for protein purification.

For purification, the supernatant was loaded onto HisTrap FF Ni-Sepharose column (Cytiva) which was pre-equilibrated with the lysis buffer. After loading the protein, the column was extensively washed with 10 and 60 mM Imidazole. The His-tagged EcoNrdR was eluted with the elution buffer (0.05 M Tris-Cl (pH 8.5 at 4°C), 0.3 M NaCl, 0.5 M Imidazole and 2 mM DTT). The purified protein was then desalted using HiPrep 26/10 Desalting column (Cytiva) equilibrated with the EcoNrdR storage buffer (0.05 M Tris-Cl (pH 8.5 at 4°C), 0.3 M NaCl and 1 mM TCEP) and stored at -80°C.

### Microscale thermophoresis

Microscale thermophoresis (MST) was used to study the binding of NrdRs to the native and mutant synthetic NrdR boxes. Oligonucleotides of 57-58 bp containing each of these NrdR boxes with flanking regions of 5 nucleotides were ordered from Sigma-Aldrich. The 5’ end of sense strand of each oligonucleotide was labeled with Cy5 by the manufacturer while the antisense strand was unlabelled (Table 8). The native and mutant synthetic oligonucleotides were obtained as 100 uM solutions.

The single stranded oligonucleotides were annealed by mixing 50 and 57.5 pmoles of the labeled and unlabeled oligonucleotides, respectively, in 50 ul buffer (10 mM Tris-Cl pH 8.0 and 50 mM NaCl) using a thermoblock. The annealing program involved incubation for 5 min at 95°C followed by gradual cooling to 25°C using 140 cycles of - 0.5°C and 45 sec per cycle, resulting in 1 µM double stranded DNA. Integrity of the annealed oligonucleotides was determined by application to Mini-PROTEAN native 5% polyacrylamide TBE gel (Biorad), staining with SYBR safe DNA gel stain (Thermo Fischer Scientific) and analysis using gel imager system C600 (Azure Biosciences) at wavelengths suitable for detection of Cy5 and Cy3 emission signal (Supplementary figure 9).

MST for analyzing the binding of EcoNrdR and ScoNrdR to the native and mutated Synt DNA fragments was performed using Monolith NT.Automated (Nanotemper Technologies, Germany). A 16-step dilution series was prepared by adding 10 μL buffer to 15 tubes. NrdR, 20 μL was placed in the first tube, and 10 μL was transferred to the second tube and mixed well by pipetting (1:1 dilution series). To each tube of the dilution series 10 μL of the binding partner (double-stranded Cy5 labelled oligonucleotide in the same buffer) was added to reach a final concentration of 10 nM. The samples were incubated for 10 min at room temperature before being transferred to Monolith™ NT.115 Series Capillaries (NanoTemper). For EcoNrdR the buffer was 0.025 M Tris-Cl (pH 8.5 at 4 °C), 100 mM NaCl, 7 mM MgCl_2_, 1 mM DTT, 0.025% Tween20, 1 mM ATP and 1 mM dATP. MST was done at high power with 50% nano-red. For ScoNrdR, the buffer was 0.025 M Tris-Cl (pH 8.0), 150 mM NaCl, 5% glycerol, 10 mM MgCl_2_, 2 mM DTT, 0.05% Tween 20, 1 mM ATP and 1 mM dATP. MST was done at medium power, 40% nano-red. The results were analysed using the MO Affinity Analysis v2.3 software (NanoTemper) with default parameters. *K*_D_ and standard deviation were calculated using fits from at least three individual titrations.

