## Supplementary Material for "Mutational and bioinformatic analysis of the binding site for the ribonucleotide reductase-specific transcriptional repressor NrdR"

### Supplementary material and comments

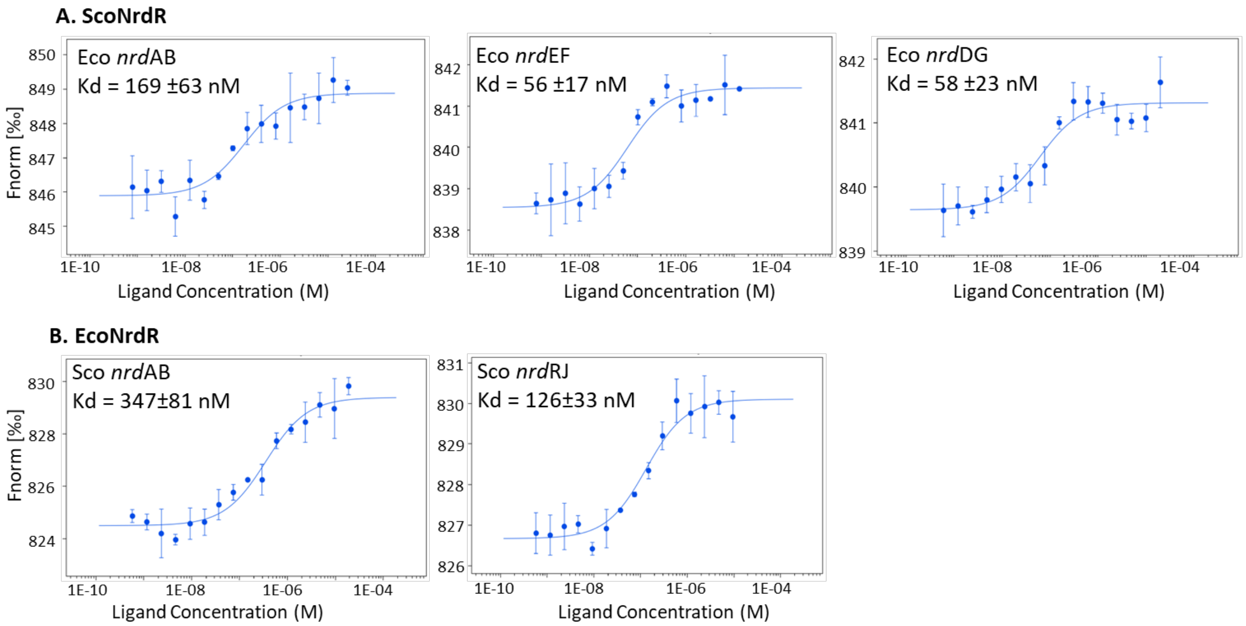


**Supplementary figure 1. Heterologous binding of NrdR to RNR promoter regions.** A. ScoNrdR binding to three *E. coli* RNR promoter regions, B. EcoNrdR binding to two *S. coelicolor* RNR promoter regions. All $K_D$ values shown in the table above.

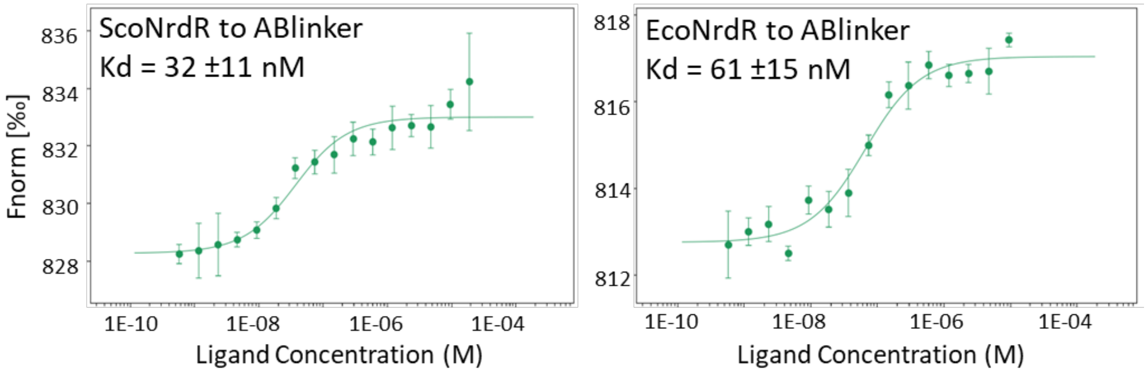


**Supplementary figure 2. Binding of ScoNrdR and EcoNrdR to the Synt DNA fragment with native *S. coelicolor nrdAB* linker region.**

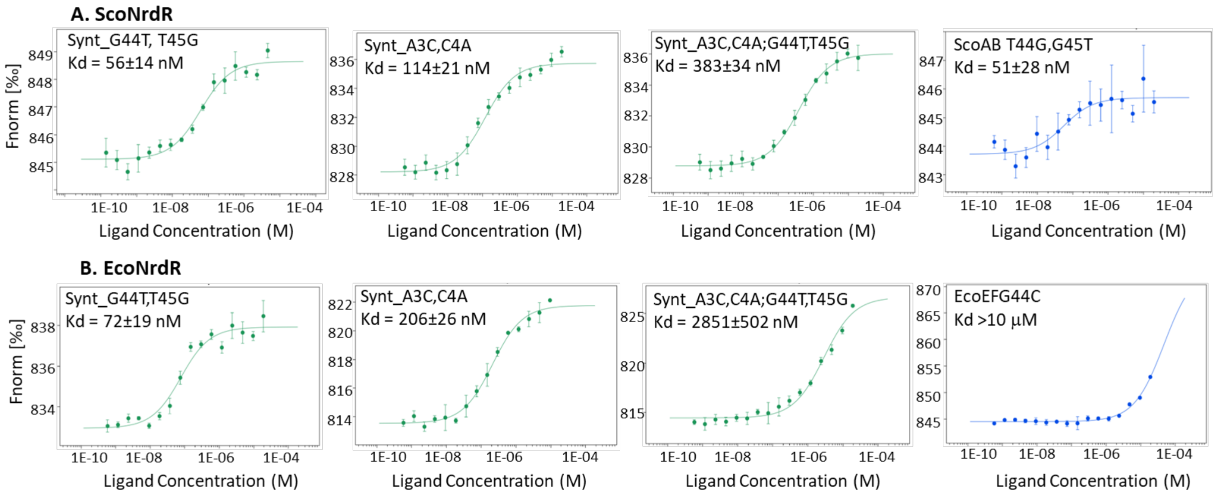


5'-gccaa**ACACAACATCTAGTGG**ttggatagcgtgagc**ACACAACATCTAGTGG**acctc-3'
4 13 35 44

**Supplementary figure 3. Binding of ScoNrdR /A) and EcoNrdR (B) to mutated Synt DNA fragments and mutated RNR promoter regions.** Green curves, binding to Synt DNA; blue curves, binding to corresponding mutations in homologous *S. coelicolor nrdAB* and *E. coli nrdEF* promoter regions, respectively. Lower line shows the sequence of a DNA fragment used with NrdR boxes in bold and mutated bp in red.

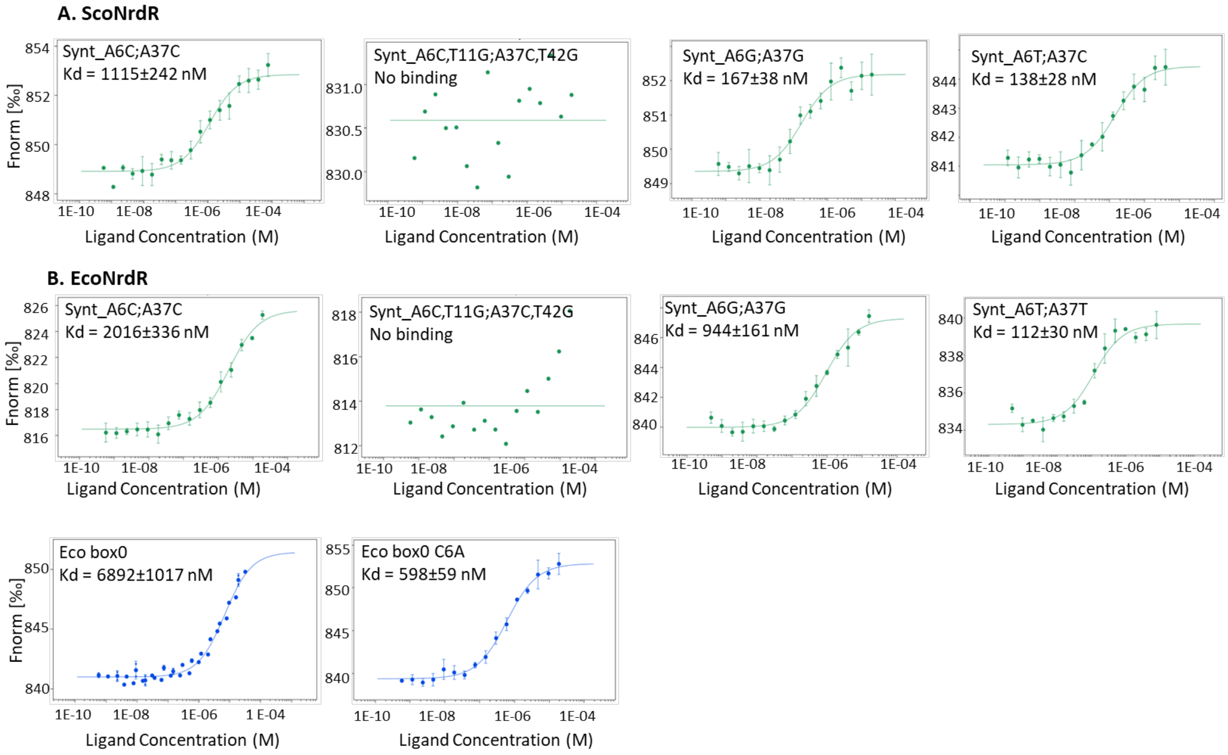


5'-gccaa**ACACAACATCTAGTGG**ttggatagcgtgagc**ACACAACATCTAGTGG**acctc-3'
6 11 37 42

**Supplementary figure 4. Binding of ScoNrdR (A) and EcoNrdR (B) to mutated Synt DNA fragments and *E. coli* RNR promoter regions.** Green curves, binding to Synt DNA. Blue curves show EcoNrdR binding to native and mutated *E. coli nrdAB* promoter fragment comprising NrdR boxes 0 and 1. Lower line shows sequence of DNA fragment used with NrdR boxes in bold and mutated bp in red.

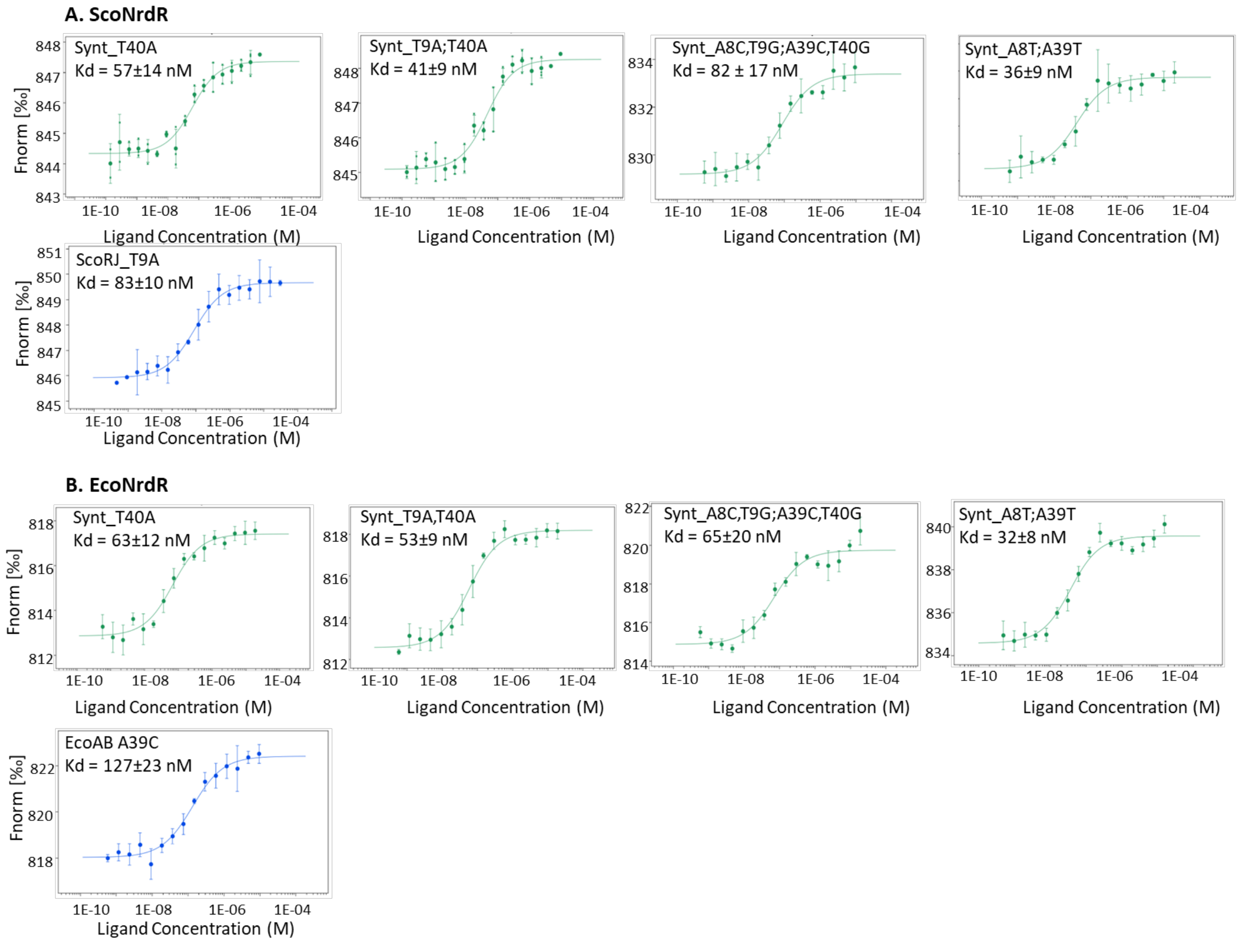


5'-gccaa**ACACAACATCTAGTGG**ttggatagcgtgagc**ACACAACATCTAGTGG**acctc-3'
8 9 39 40

**Supplementary figure 5. Binding of ScoNrdR (A) and EcoNrdR (B) to mutated Synt DNA fragments and mutated RNR promoter regions.** Green curves, binding to Synt DNA; blue curves, binding to corresponding mutations in homologous *S. coelicolor nrdRJ* and *E. coli nrdAB* promoter regions, respectively. Lower line shows sequence of DNA fragment used with NrdR boxes in bold and mutated bp in red.

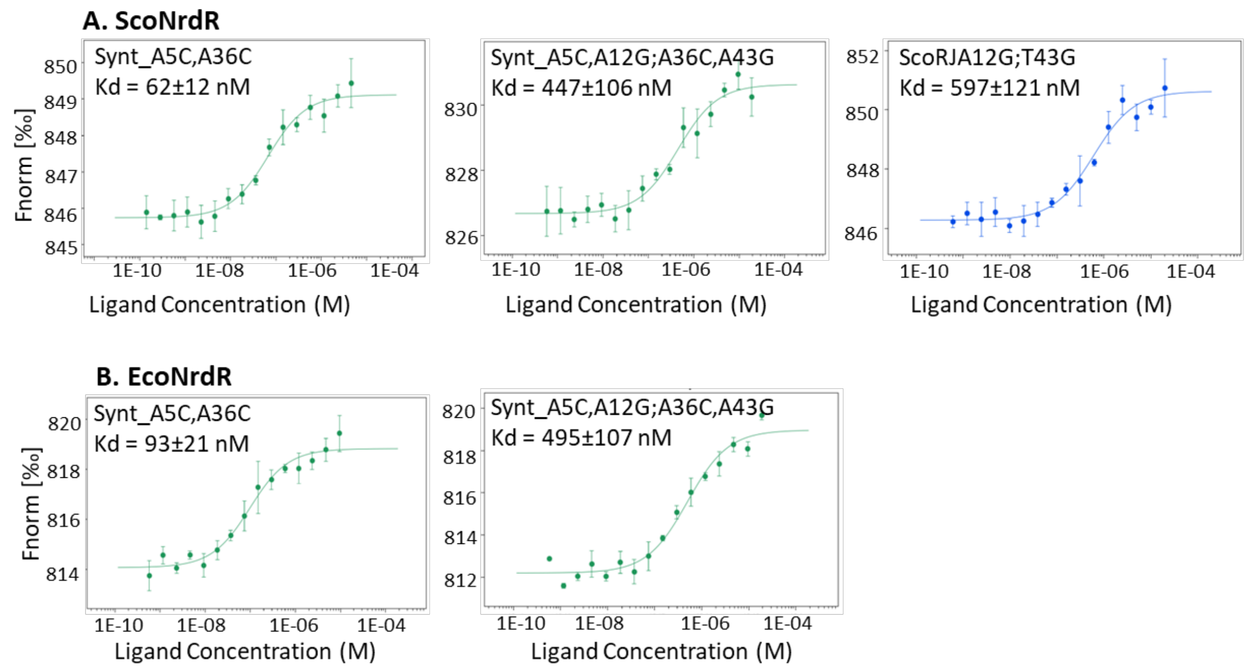


5′-gccaa**ACACAACATCTAGTGG**ttggatagcgtgagc**ACACAACATCTAGTGG**acctc-3′
5 12 36 43

**Supplementary figure 6. Binding of ScoNrdR (A) and EcoNrdR (B) to mutated Synt DNA fragments and mutated *S. coelicolor* RNR promoter regions.** Green curves, binding to Synt DNA; blue curve, binding to corresponding mutations in homologous *S. coelicolor nrdRJ* promoter region. Lower line shows sequence of DNA fragment used with NrdR boxes in bold and mutated bp in red.

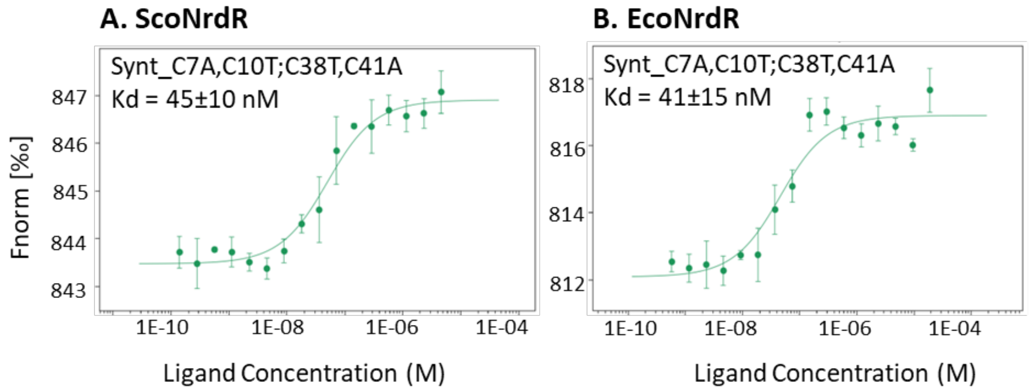


5′-gccaa**ACACAACATCTAGTGG**ttggatagcgtgagc**ACACAACATCTAGTGG**acctc-3′
7 10 38 41

**Supplementary figure 7. Binding of ScoNrdR (A) and EcoNrdR (B) to mutated Synt DNA fragments.** Green curves, binding to Synt DNA. Lower line shows the sequence of a DNA fragment used with NrdR boxes in bold and mutated bp in red.

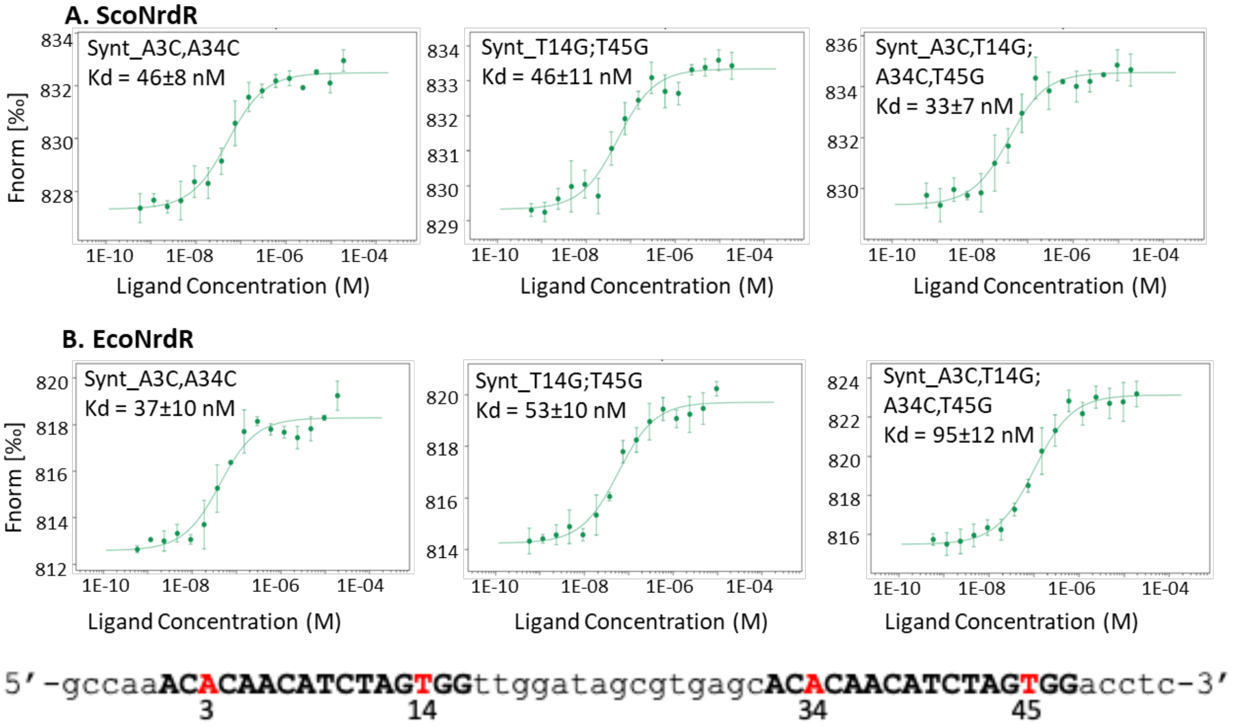


**Supplementary figure 8. Binding of ScoNrdR (A) and EcoNrdR (B) to mutated Synt DNA fragments.** Lower line shows sequence of DNA fragment used with NrdR boxes in bold and mutated bp in red.

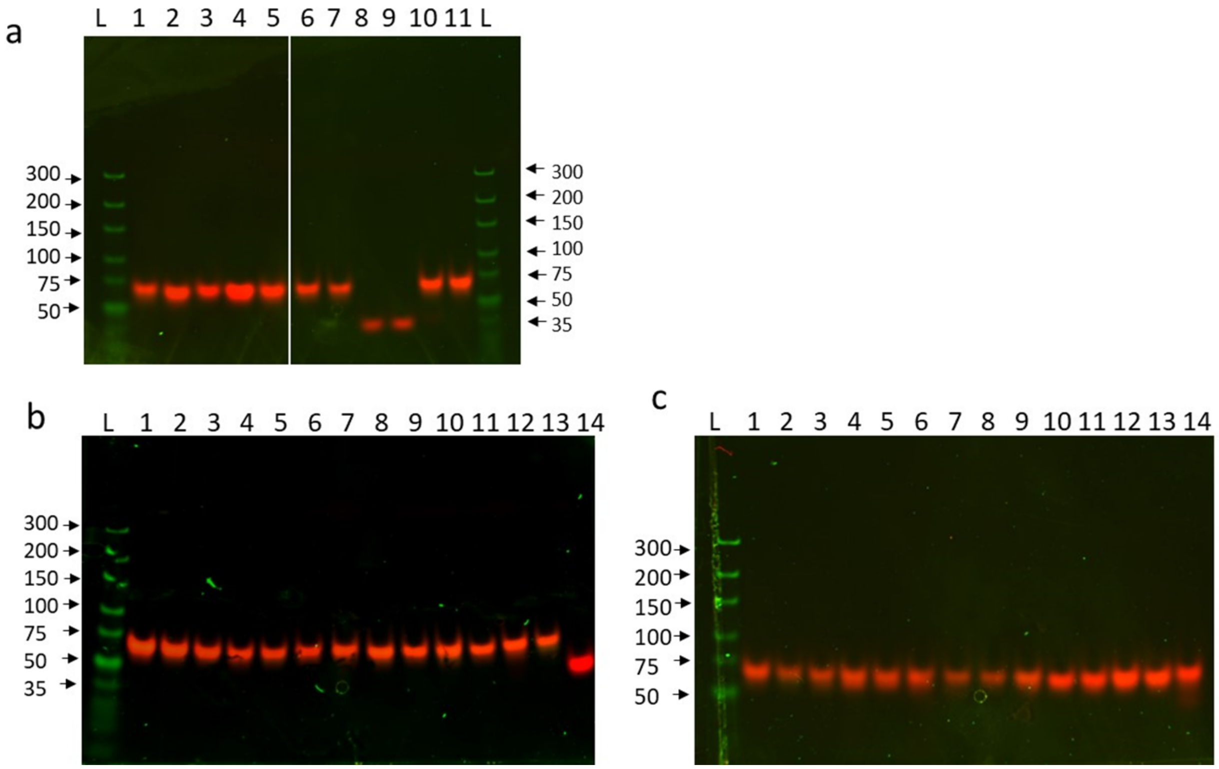


**Supplementary figure 9: PAGE of annealed oligonucleotides used for MST A:** L) DNA ladder; 1)Eco nrdAB_0,1; 2) Eco nrdAB_1,2; 3) Eco nrdEF; 4) Eco nrdDG; 5) Eco RibX; 6) Eco nrdEF dupliated box1; 7) Eco nrdEF dupliated box2; 8) Eco nrdEF box1; 9) Eco nrdEF box2; 10) Sco nrdRJ; 11) Sco nrdAB. **B:** L) DNA ladder; 1) Synt 2) Synt T9A,T40A 3) Synt T40A; 4) Synt A8C,T9G,A39C,T40G; 5) Synt A8T,A39T; 6) Synt C7T,C10A,C38T,C41A 7) Synt A6C,A37C; 8) Synt A6C,T11G,A37C,T42G; 9) Synt A6T,A37T; 10) Synt A5C,A36C; 11) Synt A5C,A12G,A36C,A43G; 12) Synt A6T,A37T; 13) Synt ScoABlinker; 14) Synt_For. **C:** L) DNA ladder; 1) Synt A3C,C4A; 2) Synt G44T,T45G; 3) Synt A3C,C4A,G44T,T45G; 4) Synt T14G,T45G; 5) Synt A3C,A34C; 6) Synt A3C,T14G,A34C,T45G; 7) Sco nrdAB T44G G45T; 8) Sco nrdRJ T9A; 9) Sco nrdRJ A12G T43G ; 10) Eco nrdEF G44C; 11) Eco nrdEF T12G G13T; 12) Eco nrdAB0,1 C6A; 13) Eco AB1,2 A39C; 14) Synt. Double stranded oligonucleotide (1 pmol) was loaded on 5% polyacrylamide gel in Tris/Borate/EDTA (TBE) buffer and ran at 70 V for ~90 min. Gene ruler Ultra low (Thermo Fischer Scientific) was used as DNA ladder.
